# Seasonal synchronization of sleep timing in industrial and pre-industrial societies: the regulating role of Daylight Saving Time

**DOI:** 10.1101/392035

**Authors:** José María Martín-Olalla

## Abstract

Did artificial light reshape human sleep/wake cycle? Most likely the answer is “yes”.

Did artificial light misalign the sleep/wake cycle in industrialized societies relative to the natural cycle of light and dark? For the *average* person —that is, obviating the tail of the distributions— the answer is probably not.

Sleep timing in extratropical, industrial (data from eight national time use surveys) societies and Subtropical, pre-industrial societies (data from nine locations coming from seven previous reports) with and without access to artificial light finds in winter a remarkable accommodation with sunrise time along a wide range of angular distance to the Equator (0° to 55°).

Within the two process model of sleep, results show sleep onset and sleep offset keep bound to each other by the homeostatic process, while the photoreceptive process aligns the phase of the sleep/wake cycle to winter sunrise. This results in a phase increasingly lagging with increasing latitude up to a delay of 120 min at 55° latitude.

In the summer season, the homeostatic process persists binding sleep onset to speep offset but Daylight Saving Time regulations in industrialized societies reduce the lag to 40 min at 55° latitude. Sleep timing is then stationary with latitude.

## I. INTRODUCTION

Sleeping and being awake are probably the most basic human activities entrained to Earth’s rotation period *T* = 24 h —the definition of hour— and the light/dark (LD) cycle. A sleep-independent circadian mechanism[1] links wakefulness to the photoperiod (light) and sleep to the scotoperiod (dark). A sleep-dependent homeostatic mechanism[1] limits our propensity to being awake to roughly 2*T/*3 = 16 h of continued wakefulness.

At the Equator the shares of light and dark are *T/*2 = 12 h, Earth’s obliquity *ε* = 23.5° alters them with latitude and seasons. The shares of the LD cycle and the shares of the sleep/wake cycle are then different and wakefulness must occur during the scotoperiod or sleep must occur during the photoperiod.

It is a wide understood that artificial light and industrialization has reshaped the human sleep/wake cycle[2] by altering the way humans receipt light during the scotoperiod. To which extend is still a matter of discussion. Much of the contemporary concern is focused in one issue: sleep deprivation caused by artificial light[3].

In the past few years several studies have addressed the sleep timing in societies with different degrees of industrialization, trying to find clues which help understanding the way modern artificial light has altered sleep timing.

Three of these studies analyzed hunter-gatherer, horticulturalist, pre-industrial societies[4–6] and reported similar or shorter sleep durations compared with industrial societies. In contrast another group of four studies reported data on homogeneous pre-industrial communities with and without access to electricity[7–10] and reported later sleep onset times and shorter sleep duration with increasing urbanization.

This manuscript will not address the impact of artificial light on sleep duration. Instead it will discuss the sources of synchronization in sleep timing. Which quantities related to the sleep/wake cycle remain stationary and which not. The study will analyze a rather eclectic collection of reports that covers industrialized countries —data coming from eight national time use surveys— and pre-industrial societies —data coming from the seven previously cited references. For so doing sleep timing and the LD cycle will be contrasted. The analysis will address the seasonal characterization of sleep timing, the dependence of sleep timing on latitude and the role of Daylight Saving Time (DST) in synchronizing sleep timing.

### II. METHODS

#### II.1. Responses and predictors

The goal of this work is a comprehensive analysis of the relation between sleep timing —sleep offset *t*_1_; the midpoint *t*_2_; and sleep onset *t*_3_— and the LD cycle.

The dependent variable, outcome or response in this analysis will be human sleep timing. It will be parameterized by the distance to solar noon, which can be evaluated from local time measurements if longitude and local time zone are known:

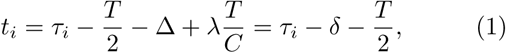

where *τ_i_* is a local time (relative to midnight), Δ is the local time zone offset relative to UTC (Coordinated Universal Time) and *λ* is local longitude relative to UTC prime meridian and *C* = 360° is one cycle. The time offset *δ* is the time distance from solar noon to midday.

The LD cycle will be characterized by the shortest photoperiod, which is that of the winter solstice. In this work “winter” or “winter solstice” will always be used in absolute form: it can indistinctly mean the Northern Solstice (June) or the Southern solstice (December) depending on the hemisphere. Calendar references will not be used.

The shortest photoperiod is the time distance from winter sunrise (WSR) to winter sunset (WSS): the time distance from the latest sunrise to the earliest sunset. It is a proxy for absolute latitude |*ɸ*|, the angular distance to Equator, irrespective of the hemisphere, to which is related by:

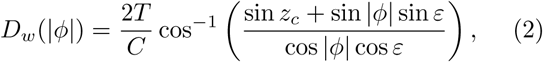

where *z*_*c*_ = −0.83° is a critical solar altitude relative to the horizon, which defines sunrise and sunset. Winter photoperiod *D*_*w*_(|*ɸ*|) is only defined and non-zero below polar circles. Above polar circles *D*_*w*_ is just set to zero (permanent scotoperiod).

The shortest photoperiod is straightforwardly related to winter sunrise and winter sunset times by:

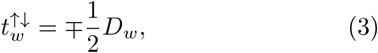

where time is given as a distance to solar noon. This simple relation suffices to understand the role of the shortest photoperiod as a predictor or independent variable in the forthcoming analysis.

Except for minor corrections due to finite angular size of the solar disk and to atmospheric refraction, the shortest scotoperiod equals the shortest photoperiod. Summer sunrise (SSR) and sunset (SSS) times are then given by ±*D_w_/*2, if time is given as a distance to midnight. Therefore the limiting boundaries of the LD cycle are characterized by a unique gradient *β* = 1*/*2 = 30 min h^*−*1^ if *D*_*w*_is the predictor.

The gradient reads thirty minute in advance/delay of winter/summer sunrise/sunset per hour change in *D*_*w*_. Events that delay with increasing distance to the Equator —winter sunrise or summer sunset— take the negative value *m* = -*β*. Events that advance with increasing distance to the Equator —winter sunset and summer sunrise— take the positive value *m* = +*β*.

Unlike boundaries, the midpoints of the LD cycle are characterized by stationary values (*m* = 0). Solar noon happens at *t* = 0, irrespective of latitude. Yearly averaged values of sunrise times and of sunset times yield values close to *∓*6 h, irrespective of latitude.

### II.2. Data sets

Table I lists the societies whose sleep timing will be analyzed in this work. It also lists geographical data relevant for the analysis and the sample size of each study. Figure 1 displays the societies on Earth’s surface. Since it is the angular distance to the Equator, and not latitude, that matters in the analysis, Figure 1 displays the surface of the Earth a unique hemisphere: Southern masses of land —blueish ink— look like upside-down.

**Figure 1.**
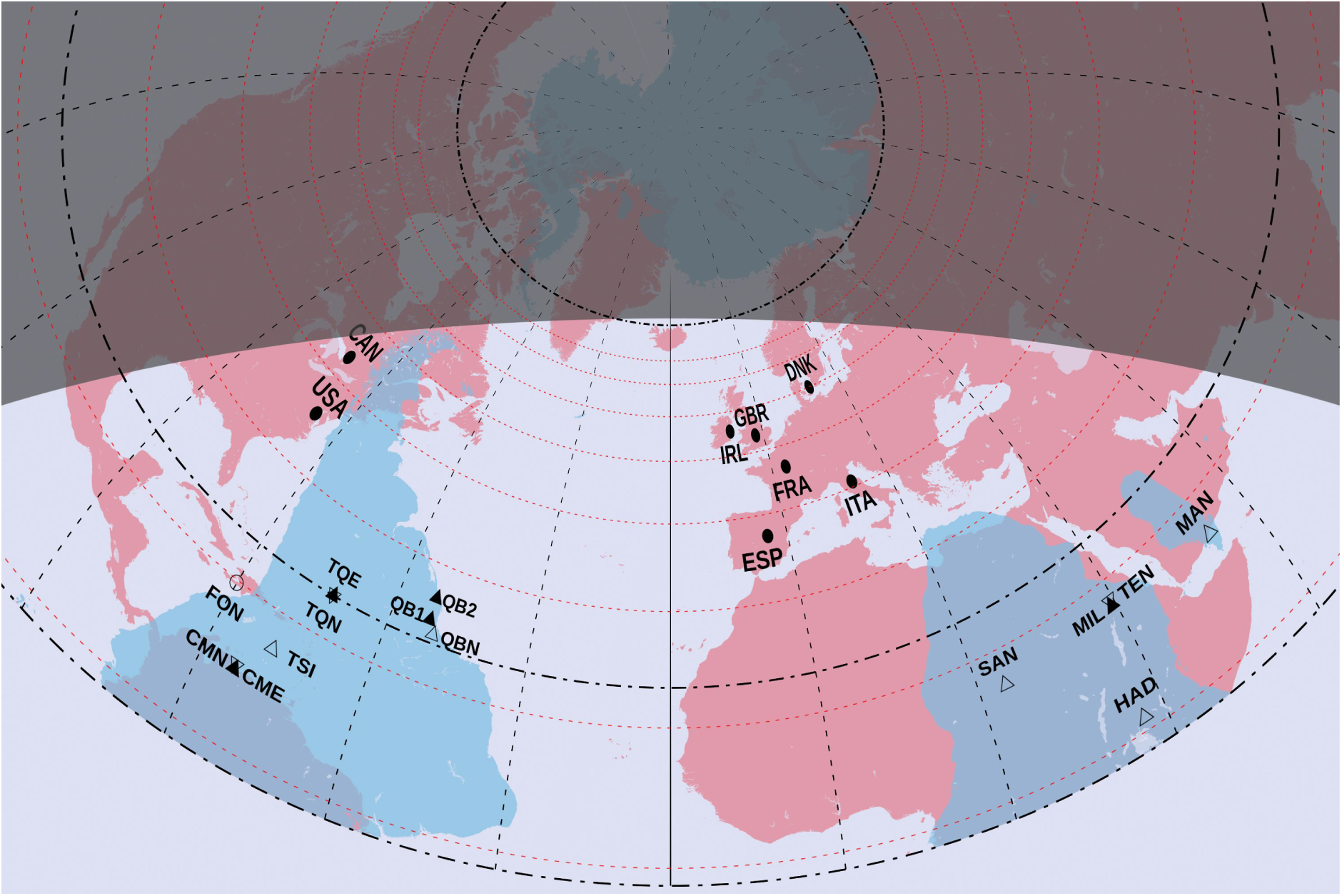
Locations of the societies whose sleep timing is analyzed in this manuscript, see Table I to decode labels. The figure shows the Earth on a unique hemisphere from Equator (bottom) to the Pole (top). The projection is azimuthal centered at *λ*_0_ = 18°8′ west of UTC prime meridian and 52° latitude. Solid symbols display societies with access to electricity. Open symbols display societies without access to electricity. Rounded symbols refer to societies in the Northern Hemisphere. Triangled symbols refer to societies in the Southern Hemisphere. Circles of latitude in heavy dash-dotted lines are the Equator, the Tropic at *ɸ* = *ε* and the Polar Circle at *ɸ* = 90° − ε. Circles of latitude in light dashed lines run values of the shortest photoperiod *D*_*w*_, increasing in steps of 1 h. They start at *D*_*w*_ = 5 h, the closest circle to the Polar Circle. The picture separates light and dark in the winter day at noon over the meridian *λ*_0_. Continental and lake polygons were taken from http://www.naturalearthdata.com/.

Societies are grouped in three categories: industrial, pre-industrial without access to electricity and preindustrial with access to electricity. Incidentally, all industrial societies lie in the Northern Hemisphere. All pre-industrial societies lie in the Southern Hemisphere, except Fondwa (FON).

Table II lists sleep timing in industrial and preindustrial societies in the winter season. Table III lists summer values.

Sleep timing in industrial societies was computed from time use surveys. They are a fundamental tool for understanding social behavior and try to ascertain when people do basic activities like sleeping, eating or working in a standard day.

This work will study results from eight national time use surveys. Six are located in Europe: Denmark[11], Ireland[12], United Kingdom[13], France[14], Italy[15] and Spain[16]. Two are located in America: Canada[17] and United States[18]. Countries are labeled according to iso-3166 alpha-3 codes.

A time survey usually takes 144 time-slots in a day, each representing ten minutes. Respondents are asked to indicate their main activity in every time-slot. The daily rhythm of an activity is the number of respondents which reported doing an activity at a given time-slot scaled by the number of respondents (see Ref. [19] for a detailed description). Therefore a daily rhythm averages the myriad of individual decisions that shapes a society.

The daily rhythm of the sleep/wake activity looks like a smoothed rectangular function with the vast majority of respondents awaken by noon and sleeping by midnight. Therefore it shows the preference of contemporary societies for monophasic sleep, only slightly modified in Spain and Italy by the siesta (nap), a short sleep at mid afternoon.

Two transitions —one in the morning, one at night (see Figure 1 in Ref. [19])— from the sleep to wake state and vice-versa are observed. A threshold located at half the range of the daily rhythm —which is close to 50 %— will help identifying sleep offset and sleep onset. They can be viewed as an average of individual wake up times and bedtimes. The average of sleep onset and sleep offset will define the midpoint of sleep (or wake).

Number of respondents in a time survey is usually in the range of thousands and differ in age, sex and social issues. Here sleep timing was computed for laborers in a weekday (Monday to Friday) and non-working population age 25 year in a week day. Table II and Table III lists the average value of this quantities.

Time use surveys usually collect data during at least one calendar year. Only the Danish and Irish surveys did not do so. Yet, there are not significant differences for the winter (here meaning autumn and winter) and summer (here meaning spring and summer) season. Modern societies entrain to clock time. Hence seasonal variations in extratropical, industrial societies are mostly induced by Daylight Saving Time (DST) onset and offset.[**?**]

Previous works on pre-industrial societies include Knutson[4] which reported sleep patterns in fifty-eight persons living in the community of Fondwa in Haiti. They have no no access to electricity. The study was conducted in January-February (winter) of 2003 using wrist actimetry. Table II and Table III collect data contained in the abstract of the manuscript, which also reports a standard deviation of 40 min.

Yetish et al.[5] reported the sleep timing in three hunter-gatherer, horticulturalist societies: the San (SAN) in Namibia, the Tsimané (TSI) in Bolivia and the Hadza (HAD) in Tanzania. None of them has access to electricity. Table II and Table III collect sleep timing listed in Table S2 of the reference, which includes winter and summer data for the San and Tsimané. Authors reported standard deviations range from 0.5 h to 1 h.

Samson et al.[6] studied sleep timing in a small-scale agricultural society named Mandena (MAN) in Madagascar. The study was conducted from July to August (winter). Table II and Table III collect data reported in Table 1 of the cited reference. Characteristically this society exhibited fragmented sleep at night but reported standard deviation amounts to 3.5 h for sleep onset and only 1 h for sleep offset.

Moreno et al.[7] analyzed sleep timing from rubber tappers in the Chico Mendes Amazon Extractive Reserve (Acre, Brazil). The study was conducted from September to November (spring) on a community without access to electricity (CMN) and a community with access to electricity (CME). Table II and Table III collect workday values of sleep timing obtained from Figure 1 of the cited reference which reports standard errors of mean values in the range of 10 min.

De la Iglesia et al.[8] analyzed sleep timing in two communities of the Toba/Qom —one with access to electricity (TQE), the other one without access (TQN)— in summer and winter. They live in the province of Formosa, the most Equatorial in Argentina. Table II and Table III collect rise times and bedtimes from Table 1 of the reference which reports standard error of the mean in the range of 10 min.

Beale et al.[9] described sleep timing in two neighboring communities in Mozambique: Milange (MIL, urban) and Tengua (TEN, rural). The study was conducted from April to June (autumn-winter). Table II and Table III collect sleep timing from Table 1 of the cited reference, which reports standard errors of the mean also in the range of 10 min.

Finally, Pilz et al.[10] investigated sleep patterns in the Quilombolas along a series of seven communities in Southern Brazil (states of Paraná, Rio Grande do Sul and São Paulo) with different degrees of urbanization. The study was conducted from March 2012 to March 2017 and Table II collects sleep timing presented in Table S2 (winter, actimetry data) of the cited reference, which reports standard deviations in the range from 30 min to 120 min. The seven locations have been grouped according to: locations with no access to electricity or which gained access only recently (two, Bombas and Areia Branca, label QBN); locations with gained access to electricity less than twenty years ago (three, São Roque, Córrego do Franco and Mamãs, label QB1); and locations with electricity for more than twenty years (two, Morro do Fortunato and Peixoto dos Botinhas, label QB2).

Only two studies[5, 8] reported data in winter and in summer. Four of the five remaining studies report winter-autumn data, and only Moreno et al.[7] reported data in spring-summer only. Except for the Quilombolas[10] —who live at 30° where the variation of sunrise times climbs close to two hours (see Table I)— data will be used indistinctly in winter and summer analysis.

There are huge methodological differences from time use surveys to studies in pre-industrial societies which must be marked as limitations. Time use surveys are large scale studies with thousands of individuals living on large area and which are entrained to standardized time. These studies are, of necessity, retrospective. On the other hand pre-industrial studies involve tens to hundreds of individuals, living on a given location. Six out of seven studies are prospective; only data from the Chico Mendes extractive reserve are retrospective.

## III. RESULTS

This section will put forward the seasonal differences in sleep timing and will contrast sleep timing against the natural gradients *m* = 0 and *m* = ±*β* through three different methods: first a visualization of data (Figure 2); then multiple binning analyses (Figure 3); and finally univariate analyses (which are included in Table II and Table III) and multiple bivariate analyses (Table IV).

**Figure 2.**
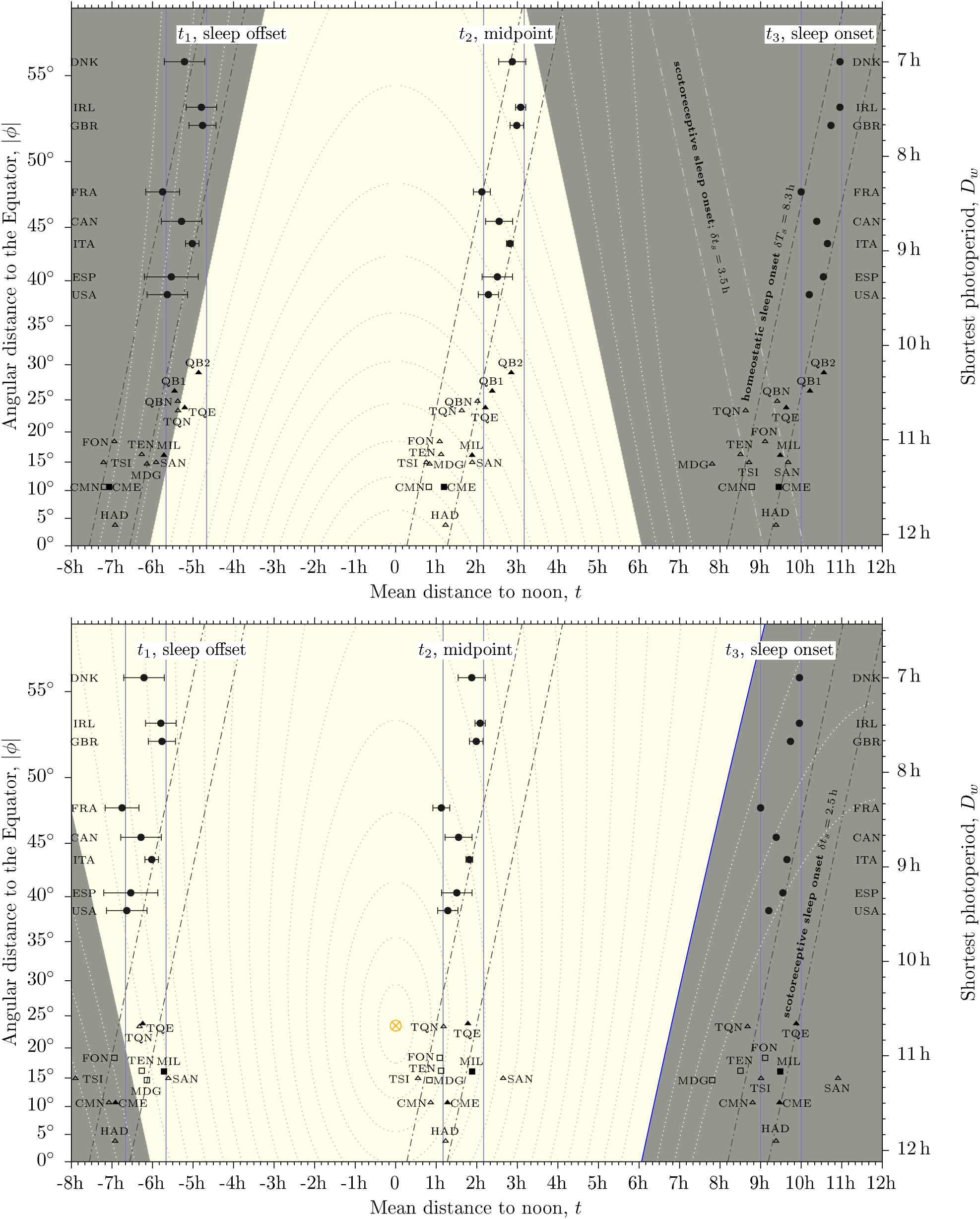
Sleep timing against shortest photoperiod in industrial and pre-industrial societies; top panel refers to winter, bottom panel to summer. Horizontal axis displays responses: sleep timing as a distance to solar noon. Vertical axes display predictors: shortest photoperiod (right) and latitude (left). Lighter background is the photoperiod; darker background is the scotoperiod. Dotted lines display solar altitude starting at *z* = −18° (outer most) in steps of 6°. The subsolar point (*z* = 90°) is noted by a crossed circle in the bottom panel. Human sleep timing is noted by open symbols (no access to electricity) and solid symbols (access to electricity). Industrial values are noted by circles; pre-industrial values by triangles. Squares are used for pre-industrial values obtained in the opposite season (summer values in top panel or winter values in bottom panel). Horizontal error lines display sleep timing from laborers in a week day (earliest bound) to standard population in week-end (latest bound). Solid, vertical (*m* = 0) lines highlight distance to noon; slanted (*m* = −*β*), dash-dotted lines, distance to winter sunrise or summer sunset; slanted dash-dot-dotted lines, distance to winter sunset or summer sunrise (*m* = +*β*). Dash-dotted lines keep still in top and bottom panels to visualize the seasonality of industrial data, which moves one hour to the left induced by DST rules. The crossed circle in bottom panel is the subsolar point, located at noon and at the Tropic *ɸ_s_* = *ε* and where Sun is overhead.

**Figure 3.**
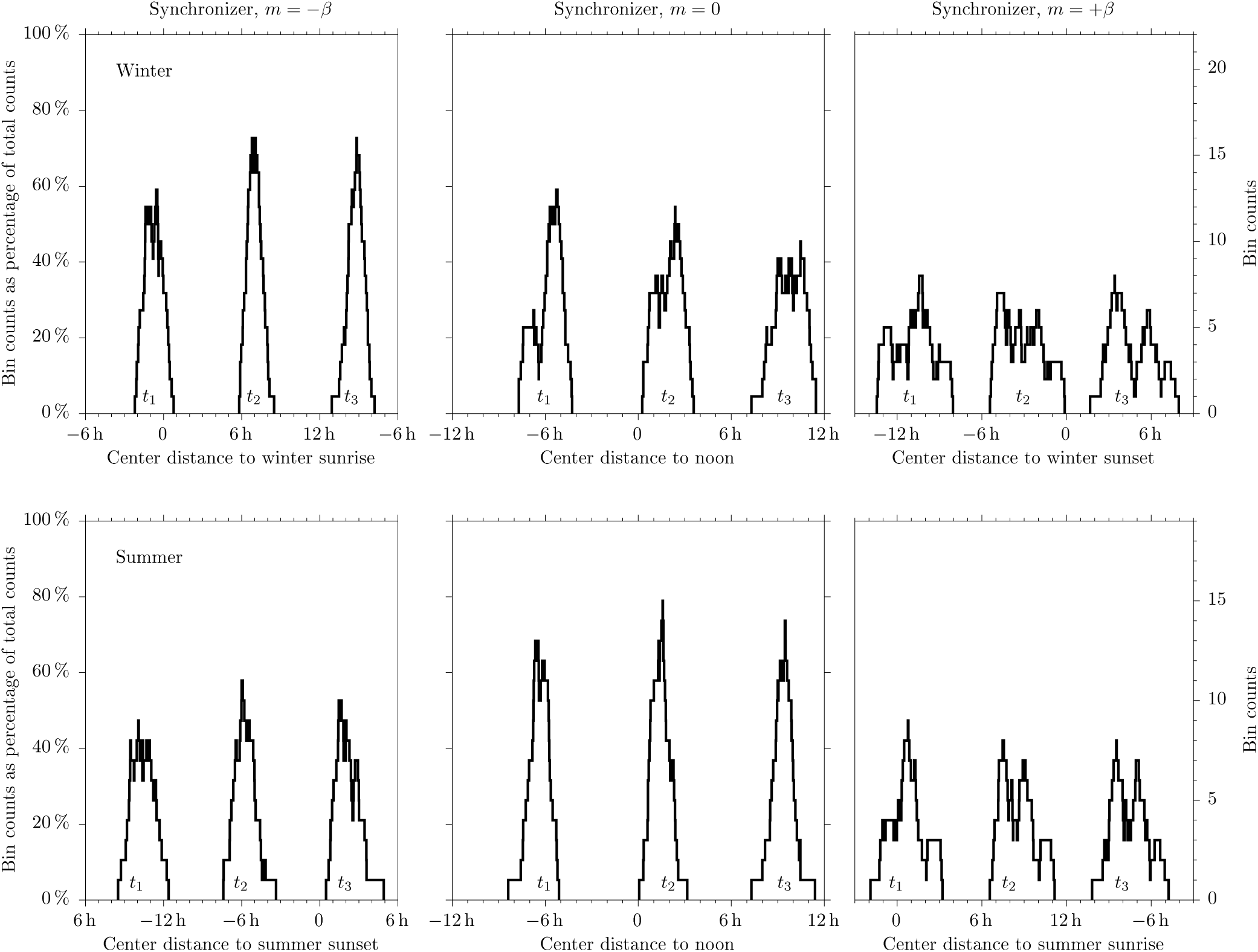
Bin counts for one-hour width bin whose center sweeps time. Top panels refer winter; bottom panels, to summer. For left to right gradient *m* = −*β* (parallel to the winter sunrise line); gradient *m* = 0 (parallel to the noon line) and gradient *m* = +*β* (parallel to the winter sunset line). Results are presented as a percentage (left axis) or as counts (right axis). Horizontal axes show time distance

Figure 2 shows human sleep timing for the winter season (top panel) and the summer season (bottom panel) against the shortest photoperiod. The figure shows sleep timing in the horizontal axis and the shortest photoperiod in the vertical axis. It is a geographical representation which mimics Figure 1. The vertical axis is latitude but displays straight values of *D*_*w*_ (right axis). The horizontal axis is longitude —distance to noon, see Equation (1)—. That way the curvilinear line of shadow in Figure 1 becomes straight (see Equation (3)). However, notice that geometrically the predictor *x* = *D*_*w*_ is displayed on the vertical axis, while responses *y* = {*t*_1_; *t*_2_; *t*_3_} are displayed on the horizontal axis. Therefore, gradients will be described relative to the upright (*m* = 0).

Each panel of Figure 2 must be understood as a three layer picture. The deepest layer shows the LD cycle: the lighter background is the photoperiod; the darker background is the scotoperiod. Dotted lines display solar altitude starting at −18° (outer most) in steps of 6°.

The middle layer is human sleep timing. Solid symbols refer to people with access to electricity; open symbols, to people without access to electricity. Circles display industrial values; triangles pre-industrial societies; squares in top panel refers to data obtained in the summer season; squares in bottom panel refers to data obtained in the winter season.

Horizontal error lines in industrial data span from sleep timing for laborers in a week day (earliest bound) to sleep timing for non-laborers in a week day aged 25 years or more (latest bound). Error lines for sleep onset are too short to be displayed.

The aboveboard layer shows straight lines with gradients *m* = −*β* (dash-dot), *m* = 0 (solid) and *m* = +*β* (dash-dot-dot). They represent the natural gradients associated to the boundaries of the LD cycle (*m* = ±*β*) and to noon (*m* = 0).

Adjacent straight lines in Figure 2 are always separated by one whole hour. They bin data points within the interval of one hour. Bin position is uniquely determined by the center of the bin. Bins can move along the horizontal axis and the number of counts inside the bin can be recorded.

This technique can be used to put forward which gradient best collects data. Results are shown in Figure 3. In winter (top panels) gradient *m* = −*β* gets the narrowest and highest peak, positioning sleep timing one hour before sunrise (offset), seven hours after sunrise (midpoint) and eight hours before sunrise (onset). Gradient *m* = 0 displays a high peak for sleep offset but it is wider than peak for *m* = −*β*. Gradient *m* = +*β* poorly interact with data.

In summer (bottom panels) it is *m* = 0 which exhibits the best coherence. Sleep offset is located six hours before noon, the midpoint is two hours after noon and sleep onset occurs nine hours after noon. Gradients *m* = −*β* and *m* = +*β* display lower and smeared peaks. Nonetheless, sleep onset is located some two hours after sunset and sleep offset is locate around sunrise.

Graphical information shown in Figure 3 can also be described by the univariate analysis of sleep timing listed in Table II (winter) and Table III (summer). The point to note is the variability. In winter, variability decreases by if sleep timing is measured relative to sunrise *t_i_ − βD_w_*, compared to measurements relative to noon *t*_*i*_. In summer, it is just the opposite. Values shown in blue ink highlight this change over.

Finally Table IV shows bivariate analysis —multilinear regression analysis— in which *D*_*w*_ is the predictor and sleep timing is the response. It should be stressed that no regression lines are displayed on Figure 2 which, for the sake of simplicity, only shows lines whose gradient is related to the LD cycle.

Results in Table IV are shown line by line. They include: (1) sample gradient *m*_*s*_ and (2) its confidence interval *CI* both measured in units of *β* = 30 min h^*−*1^, except when the response is solar altitude, in which case they are measured in degrees per hour, the parentheses show a symmetrical semi-interval of confidence around *m*_*s*_; (3) the probabilistic value *p*; (4) the Pearson’s coefficient; and (5) possible outliers —experimental data whose confidence residual excludes zero. Confidence is always taken at the level *α* = 5 %. Probabilistic values smaller than this level are highlighted in blue ink along with the corresponding Pearson’s coefficient. In these cases the null hypothesis “response does not depend on the predictor” is rejected. Gradients and confidence intervals for the regression relative to winter sunrise are not displayed: they can be obtained just by adding 1 to numerical values shown for the regression relative to noon.

For the winter season sleep timing gets a gradient close to *β*: *m*_*s*_ = −0.77(37); −0.85(28); −0.95(35) *β*. Pearson’s coefficient are *R*^2^ = 0.49; 0.66; 0.62 and probabilistic values are smaller that *α*. Therefore the null hypothesis “sleep timing —offset, midpoint and onset— distance to noon does not depend on the shortest photoperiod” is rejected in winter. That means sleep timing depends on the predictor. Taking into account that the range of the predictor is 300 min (see Table I), the predicted lag in the phase of the sleep/wake cycle from the Equator to |*ɸ*| = 55° amounts to *m*_*s*_ × Δ*D*_*w*_ ~ 120 min or two hours.

However, if sleep timing is measured relative to winter sunrise then the probabilistic values are larger than the confidence level. Therefore the null hypothesis “sleep timing —offset, midpoint, and onset— distance to winter sunrise does not depend on the shortest photoperiod” sustains at the confidence level.

Also interestingly, winter solar altitude at sleep offset sustains the null hypothesis (*p* = 0.73), which means winter solar altitude at sleep offset is stationary with the shortest photoperiod. The average value *z*_*w*_ ~ −9° (see Table II) is located in the middle of the winter nautical twilight.

Still in winter sleep onset to sleep offset *t*_3_ − *t*_1_ − *T* hits a gradient *m*_*s*_ = −0.18(43)*β* with probabilistic value *p* = 0.38 larger than the confidence level. Therefore the null hypothesis “sleep onset to sleep offset does not depend on the shortest photoperiod” sustains at the confidence level. This result comes from the fact that sleep offset and sleep onset follow similar trends: the difference offsets the trend.

As far as the summer season is concerned the conclusions are again reversed: the null hypothesis “sleep timing distance to noon does not depend on the shortest photoperiod” sustains at the confidence level with probabilistic values *p* = {0.11; 0.07; 0.16}. Sample gradients *m*_*s*_ are one third smaller than those reported in the winter season and hit *m*_*s*_ ~ −0.28*β* ~ −8 min h^*−*1^. Now for the observed range of the predictor the lag in the phase of the sleep/wake cycle amounts to *m*_*s*_ × Δ*D*_*w*_ ~ 40 min, compared to 120 min in winter.

## IV. DISCUSSION

The two process model of sleep[1, 20, 21] describes the sleep/wake cycle with three gold standards: (1) the circadian regulation of sleep (process C) creates a photoreceptive propensity for wakefulness during the day; therefore people tend to abhor waking up before sunrise; (2) it also creates a scotoreceptive propensity for sleep at night; hence people tend to abhor going to bed before sunset. Likewise, (3) the homeostatic regulation of sleep (process S) creates a propensity for sleep with increasing duration of wakefulness, which generally saturates (and triggers sleep) when duration is close to 2*T/*3 = 16 h. All three standards meet in Figure 2 within the range of observed latitudes.

In winter (see top panel) the first process photoreceptively aligns sleep offset to sunrise —the ancient time reference—, instead of noon —the modern time reference—. Bivariate analysis (see Table IV) reports stationary values of winter solar altitude at winter sleep offset, and stationary values of sleep offset distance to winter sunrise. They are a supporting evidence for the photoreceptive regulation of winter sleep offset along a wide range of latitudes. Below *D*_*w*_ ~ 7 h (or above |*ɸ*| ~ 55°) winter sunrise is so delayed that people may wake up long before it, induced by process S. Evidences based in the “sleep and other personal care” cycle suggest[19] this cross-over.

A scotoreceptive (photoreceptive) circadian (C) regulation of sleep onset in figure 2 (top panel) can only be dictated by the gradient *m* = +*β*, shown by dash-dot-dotted lines in the figure, whose distance to winter sunset is stationary. This gradient poorly interact with data (see Figure 3 top, right). However pre-industrial societies without electricity (open triangles in Figure 2) can be binned inside a one-hour width bin delayed *δt_s_* = 3.5 h from winter sunset. In this category only Mandena sleep onset is not binned as a result of their preference for segmented sleep. Toba/Qom, Milange/Tengua and Chico Mendes locations, which pair societies with and without access to electricity find the member without electricity on the near side of the photoreceptive bin, and the member with electricity on the far side of the bin. The Quilombola location without electricity meets the far side of the bin but those with electricity lie well outside the bin.

A homeostatic regulation (S) of sleep onset can only be dictated by the synchronizer of sleep offset (*m* = −*β*, dash-dot lines in Figure 2). Sleep onset in industrial societies can be homeostaticly binned with center at *δT* = 8.3 h to next sleep offset or 15.7 h from previous sleep offset. This behavior differs from the synchronization of working activity, whose start times (morning) syncs to the photoreceptive (*m* = −*β*) mechanism but whose end times (evening) again happens to be regulated by a photoreceptive mechanism, which now is *m* = +*β*.[19]

As observed in Figure 2 (top) the photoreceptive bin (*m* = +*β*) overlaps with the homeostatic bin (*m* = −*β*) at *D*_*w*_ ~ 11.25 h. Interestingly pre-industrial sleep onset can be indistinctly binned by both mechanisms, perhaps characterizing homeostaticly and photoreceptively natural sleep onset in winter. However at the latitude of Spain and United States the homeostatic process is delayed by 2 h; at the latitude of France it makes 3 h, and at the latitude of the British Isles makes 4 h. The rate of increase equals the rate of decrease of the shortest photoperiod.

Ancient sleep onset in Europe may have been located anywhere between these two mechanisms. The lack of efficient, cheap artificial light, the temperature and the intense physical labor would have favored the circadian, photoreceptive mechanism. However, still in that case, the photoreceptive sleep onset comes only some 11 h after sunrise in Britain and only some 13 h after sunrise in Spain. In contrast, pre-industrial societies sustain at least 14 h of continued wake (see sleep onset to sleep offset in Table II). The low value of continued wake would have played against the photoreceptive mechanism if the homeostatic process played also a role, as suggested by pre-industrial sleep timing. Very understandably, earlier sleep onset —somewhat bound to winter sunset— should have led to segmented sleep[22, 23]. Interestingly enough present day dinner times in Europe show synchronization with winter sunset times[19] and are placed within the photoreceptive strip shown in Figure 2: some three hours after winter sunset.

There are only seven pre-industrial sleep timing available in the summer season (see triangles in Figure 2 bottom panel or unstarred societies in Table III). Only four of them (the Tsimané, the San and the two Toba/Qom) allow for a comparison with winter sleep timing. Two of them (the Tsimané and the San) live at |*ɸ*| = 15° where the seasonal variation of sunrise times amounts to 50 min or 0.88 h (see Table I). Yet they exhibit markedly different behavior. The Tsimané do track solar variations: in summer they advance sleep offset and delay sleep onset by quantities smaller that 50 min. The San delay sleep timing, up to 70 min (sleep onset). This may be related to the impact of other quantities like temperature in a desertic environment[5].

The Toba/Qom live at the Tropic where the seasonal variation of sunrise times exceeds one hour (see Table I). Toba/Qom advance sleep offset by one hour in summer, irrespective of access/no access to electricity. Yet they wake up after sunrise (see Figure 2 and Table III). Sleep onset does not change (without electricity) or delays by half an hour (with electricity).

Industrial sleep timing is advanced by one hour induced by DST rules. The size of the change is similar to those discussed for pre-industrial sleep timing, but unlike them it is characteristically a change of phase: it equally advances sleep offset and sleep onset. It also should be mentioned that the variation of sunrise times amounts to 4 h on average.

With all that in mind, discussing the role of the photoreceptive circadian mechanism in summer sleep timing is complicated. Sleep onset in pre-industrial values lie in a scotoreceptive bin located *δt_s_* = 2.5 h after summer sunset. It is one hour earlier than the scotoreceptive bin observed in winter. Industrial sleep timing increasingly advance relative to this bin with increasing latitude. This may suggest that the photoreceptive circadian process is less efficient than the homeostatic process in regulating sleep timing through seasons. Notice that the photoreceptive mechanism is impacted by the seasonal variations of the LD cycle while the homeostatic mechanism is nonseasonal. Indeed, sleep onset to sleep offset is stationary with *D*_*w*_ (or |*ɸ*|) both in summer and in winter (see Table IV).

Winter regulation of sleep timing through the photoreceptive and homeostatic mechanisms creates a lag in the phase of the sleep/wake cycle which amounts to 120 min at 55°. This lag is not intended to mean necessarily a circadian misalignment. The question is: can it survive to the opposite season?

DST rules regulate the phase of the sleep/wake cycle and offset the winter lag as observed in figure 2 and noted by the values of sample gradients for winter and summer bivariate analysis: *m_s_ ~ −β* (winter) and *m_s_ ~ −β/*3 (summer), see Table IV. It was previously argued[24] that DST changes are theoretically equivalent to a geographical translocation which moves industrialized timing “towards the Equator” (from Central Germany to Morocco without changing climate as authors said in a much narrower discussion than this one). However, it is the Sun that travels *towards the observer* from winter to summer and rises 2*ε* = 47° in the sky at extratropical latitudes. This is more than twice the difference in latitude from Germany to Morocco.

The point to note in Figure 2 is the subsolar point. At the winter solstice (top panel) it lies beneath the bottom axis of the panel at |*ɸ_s_*| = 23.5°. The winter angular distance to the subsolar point is simply |*ɸ*| + |*ɸ_s_*|. It increases with increasing latitude up to a remarkable value of ~ 80° for Denmark. That means the Sun only rises 10° above horizon at noon in winter.

By the summer solstice the subsolar point has traveled enough to be placed in the bottom panel (see the yellowish crossed circle). Now the angular distance to the subsolar point is |*ɸ* − *ɸ_s_*|. It is not dictated by the distance to the Equator (latitude) but by the distance to the Tropic. As a result summer solar altitude reaches at noon is 90° − ε = 66.5° at the Equator and at |*ɸ*| = 47° as well[**?**] as lines of solar elevation angles in Figure 2 (bottom panel) reveals. Light conditions are then more homogeneous in summer than in winter. Therefore it should not be a surprise that sleep timing were also stationary.

Finally it should be discussed if either summer sleep timing could survive in winter or if winter sleep timing could survive in summer. The first option would mean permanent DST in industrialized, extratropical countries. The photoreceptive process (C) is then challenged with sleep offset occurring well before winter sunrise which breaks one of the golden rules.

Many times this is not socially welcome and the arrangement does not sustain too long as experiences in United Kingdom (1970s), Portugal (1970s and 1990s), Chile (2015) or Russia (2014) show. But, also, many times it is socially accepted: Alaska, Saskatchewan (1956), Iceland (1969) and, perhaps, Región de Magallanes in Chile (2016), all of them at *ɸ* = 50° or beyond. While culture plays a role in addressing this issue, latitude is also determining. First, as latitude increases and the winter photoperiod becomes short enough, the social propensity for waking up long before sunrise may increase, forecasting the delay in winter sunrise and the advance in winter sunset. Second, as latitude increases the rate of seasonal variations also increases, perhaps making harder to follow them. Helped by artificial light some societies at this range of latitudes are developing a non-seasonal behavior where people are just mere observers of the changes in the scotoperiod and photoperiod.

The second option would mean permanent winter time. Therefore the winter lag would sustain in the summer season. This is noted in Figure 2 (bottom panel) by the surviving dash-dot lines with gradient *m* = *−β*.

From the point of view of the two model process it seems that nothing has changed from winter to summer but it must be noted that sleep onset is now photoreceptively bound to sunset, while sleep offset is now homeostaticly bound to sleep onset. Although the arrangement looks fine enough, another accommodation is found by advancing sleep timing, thus reducing the lag inherited from winter.

The foremost evidence of this idea is the success of DST rules in many industrial, extratropical societies. Diurnal sleep is mandatory in summer at this range of circles of latitude, but the winter lag proved to be too much for the summer. The history of DST always shows some “clever” people in early 20th century trying to prevent others from resting in the summer mornings.[25] Long before that, in 1810, the Cádiz Cortes (Spanish first national assembly) seasonally advanced the timing of their sitting by one hour from May to October[**?**], a change which is equivalent to modern DST, including its transitions.

The solar altitude at noon, which is related to latitude, also influences the problem: the advance in summer sleep timing is a rational, efficient way of preventing insolation and overheating in the central hours of the summer day. This behavior is observed also in pre-industrial societies (see supplementary material in Yetish et al.[5]). It likely plays a role in understanding early sleep offset in the Tsimané, the Hadza, Chico Mendes and Fondwa.

Looking in retrospect to Subtropical, pre-industrial data and to the cycle of light and dark, one hour of seasonal variation in human activity at extratropical latitudes is not an oddity. Once time was standardized, societies got entrained to clock time and got eager for year round timing, one hour was the minimum available amount of change per coup to adapt seasonally social timing. The impact of this change on the circadian system is a concern in terms of public health[26, 27] after decades of analysis but for that to be fully addressed it is also necessary to understand the impact of a non-seasonal behavior in public health.

## V. CONCLUSIONS

Winter sleep timing in pre-industrial, hunter/gatherer-horticulturalist, Subtropical societies and in Extratropical, industrial societies remarkably accommodates to the cycle of light and dark across a wide range of angular distance to Equator (0° to 55°) and social development. Therefore, the phase of the sleep/wake cycle lags relative to noon/midnight as latitude increases up to 120 min at 55°. In summer, daylight saving time in industrial societies largely turns the phase of the sleep/wake cycle stationary with latitude.

Artificial light notably impacted the way in which the average person handles her wakefulness during the scotoperiod, after sunset. However it may not have misaligned the sleep/wake cycle since its phase is still governed by natural trends.

Daylight saving time is on public debate in Europe and America. The issue, which is influenced by latitude, is governed by a trichotomy. First option is sleep timing should sustain noon synchronization year round —which identifies to permanent summer time— and, as a result, sleep offset would occur long before sunrise at Extratropical latitudes. The second option is sleep timing should sustain winter synchronization year round —which identifies to winter time— and, as a result, the phase of the sleep/wake cycle should increasingly lag from noon with increasing latitude. This would increase the exposure in the central hours of the day at the hottest season of the year. The third option is sleep timing should seasonally swing to some extend. In this case DST is an appropriate option for contemporary societies bound to clock time, where the change of time can only proceed by the coup of one hour.

## DATA AVAILABILITY

Time Use Surveys are official surveys carried out by public Institutions. In US it is the Bureau of Labor Statistics and the United States Census Bureau. In UK it is the Office of National Statistics. They are a key tool for sociologists and economists to understand how we share time in a standard day. Any of them is a traceable, well-identified set of data.

Spanish, Dansih and American Time Use Survey microdata were freely available in the internet at the time of writing this manuscript. American Time Use Survey data can be obtained at https://www.bls.gov/tus/data.htm. A link to these surveys is provided in their corresponding references.

French, British, Italian, Canadian and Irish Time Use Survey microdata were obtained after a formal request was addressed to the institutions. The microdata were then sent to me by email, a hard-copy or as a downloadable link. In 2015, these institutions definitely were happy to send their microdata to any researcher who may need them.

Some other Institutions do not release their microdata, or request a fee for them, or request the researcher to visit the Institution if she needs access to them. In these cases I simply refused obtaining the microdata.

## ACKNOWLEDGMENTS

The author expresses his gratitude to the institutions which granted access to time use survey microdata.

Time use survey microdata came as a `csv` text file except Irish and French data. In these cases `PSPP` was used to generate a `csv` text file. The text file was parsed with a `language C` code compiled by gcc 5.4 and read in `octave 4.0`. This software was used to compute sleep timing. Its function `regress` was used for multiple linear regression analysis. All tabular material and figures were automatically produced from the same input files containing sleep timing. Manuscript was originally written in LATEX 2_*ε*_, typed in `GNU Emacs` 24.5 assisted by `AucTeX 12.1`. Graphs were produced by `gnuplot 5.1 cairolatex` terminal, except Figure 1 which was produced in the `pngcairo` terminal, xplanet took care of the azimuthal projection. `Mendeley-Desktop 1.9` helped handling bibliography. All this on three different computers each running a `Xubuntu 16.04 LTS Xenial Xerus` distro and synced by `ownCloud. Mendeley` and `ownCloud` services were locally provided by author’s institution Universidad de Sevilla.

## Appendix A: Tabular material

**Table I.**
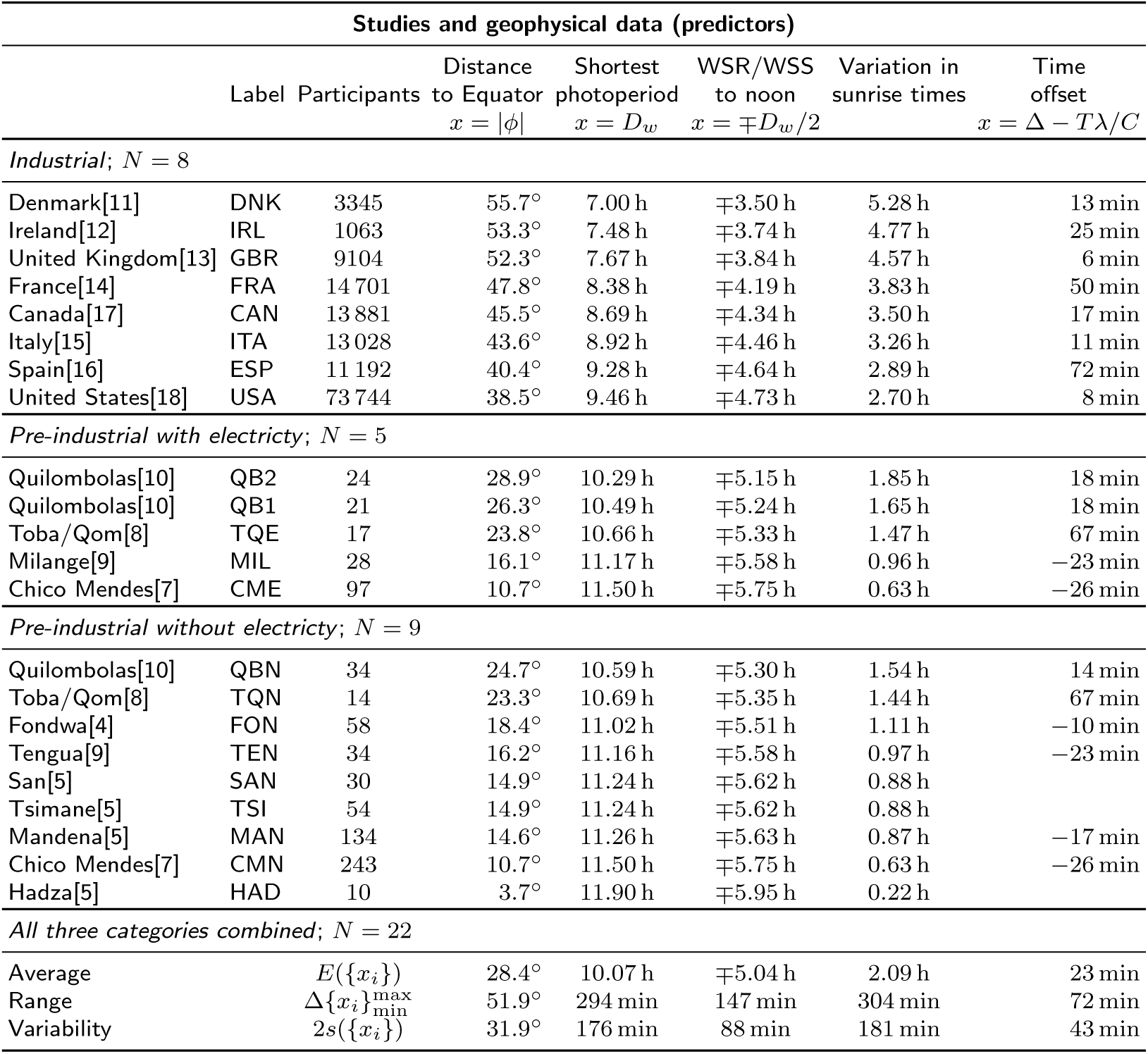
Overview of the relevant geographical information. Geographical data in industrial countries are population-weighted median values extracted from the database of cities with a population larger than 1000 inhabitants at http://www.geonames.org; labels are ISO 3166-1 alpha-3 codes. The shortest photoperiod is a function of |*ɸ*| and Earth’s obliquity. Winter sunrise WSR time and winter sunset WSS time are given as a distance to solar noon in decimal hours. The variation of sunrise times is the difference between summer rise time and winter sunrise time. Time offset is the difference between solar noon and local time midday (rounded to one minute). In summer one hour must be added in industrial societies to account for DST rules. Univariate analysis sample average, sample range and the variability expressed as twice the sample standard deviation are shown for all three categories combined. See figure 1 for a map showing all locations.

| Studies and geophysical data (predictors) |  |  |  |  |  |  |  |
| --- | --- | --- | --- | --- | --- | --- | --- |
| | Label | Participants | Distance<br>to Equator<br>$x = \vert \phi\vert $ | Shortest<br>photoperiod<br>$x = D_w$ | WSR/WSS<br>to noon<br>$x = \mp D_w/2$ | Variation in<br>sunrise times | Time<br>offset<br>$x = \Delta - T\lambda/C$ |
| *Industrial; N = 8* |  |  |  |  |  |  |  |
| Denmark[11] | DNK | 3345 | 55.7° | 7.00 h | $\mp 3.50$ h | 5.28 h | 13 min |
| Ireland[12] | IRL | 1063 | 53.3° | 7.48 h | $\mp 3.74$ h | 4.77 h | 25 min |
| United Kingdom[13] | GBR | 9104 | 52.3° | 7.67 h | $\mp 3.84$ h | 4.57 h | 6 min |
| France[14] | FRA | 14 701 | 47.8° | 8.38 h | $\mp 4.19$ h | 3.83 h | 50 min |
| Canada[17] | CAN | 13 881 | 45.5° | 8.69 h | $\mp 4.34$ h | 3.50 h | 17 min |
| Italy[15] | ITA | 13 028 | 43.6° | 8.92 h | $\mp 4.46$ h | 3.26 h | 11 min |
| Spain[16] | ESP | 11 192 | 40.4° | 9.28 h | $\mp 4.64$ h | 2.89 h | 72 min |
| United States[18] | USA | 73 744 | 38.5° | 9.46 h | $\mp 4.73$ h | 2.70 h | 8 min |
| *Pre-industrial with electricity; N = 5* |  |  |  |  |  |  |  |
| Quilombolas[10] | QB2 | 24 | 28.9° | 10.29 h | $\mp 5.15$ h | 1.85 h | 18 min |
| Quilombolas[10] | QB1 | 21 | 26.3° | 10.49 h | $\mp 5.24$ h | 1.65 h | 18 min |
| Toba/Qom[8] | TQE | 17 | 23.8° | 10.66 h | $\mp 5.33$ h | 1.47 h | 67 min |
| Milange[9] | MIL | 28 | 16.1° | 11.17 h | $\mp 5.58$ h | 0.96 h | −23 min |
| Chico Mendes[7] | CME | 97 | 10.7° | 11.50 h | $\mp 5.75$ h | 0.63 h | −26 min |
| *Pre-industrial without electricity; N = 9* |  |  |  |  |  |  |  |
| Quilombolas[10] | QBN | 34 | 24.7° | 10.59 h | $\mp 5.30$ h | 1.54 h | 14 min |
| Toba/Qom[8] | TQN | 14 | 23.3° | 10.69 h | $\mp 5.35$ h | 1.44 h | 67 min |
| Fondwa[4] | FON | 58 | 18.4° | 11.02 h | $\mp 5.51$ h | 1.11 h | −10 min |
| Tengua[9] | TEN | 34 | 16.2° | 11.16 h | $\mp 5.58$ h | 0.97 h | −23 min |
| San[5] | SAN | 30 | 14.9° | 11.24 h | $\mp 5.62$ h | 0.88 h | |
| Tsimane[5] | TSI | 54 | 14.9° | 11.24 h | $\mp 5.62$ h | 0.88 h | |
| Mandena[5] | MAN | 134 | 14.6° | 11.26 h | $\mp 5.63$ h | 0.87 h | −17 min |
| Chico Mendes[7] | CMN | 243 | 10.7° | 11.50 h | $\mp 5.75$ h | 0.63 h | −26 min |
| Hadza[5] | HAD | 10 | 3.7° | 11.90 h | $\mp 5.95$ h | 0.22 h | |
| *All three categories combined; N = 22* |  |  |  |  |  |  |  |
| Average | $E(\{x_i\})$ | | 28.4° | 10.07 h | $\mp 5.04$ h | 2.09 h | 23 min |
| Range | $\Delta\{x_i\}_{\min}^{\max}$ | | 51.9° | 294 min | 147 min | 304 min | 72 min |
| Variability | $2s(\{x_i\})$ | | 31.9° | 176 min | 88 min | 181 min | 43 min |

**Table II.** Sleep timing in industrial and in pre-industrial societies with and without electricity. Times are expressed in decimal hours as a distance to solar noon *t*_*i*_ or a distance to winter sunrise (WSR) *t*_*i*_ − *βD_w_*. In the first case, the time offset (see Table I) and *T/*2 = 12 h must be added to obtain local times. Winter solar altitude WSA at sleep offset is also shown. Finally sleep onset also lists the distance to next sleep offset. Univariate analysis for all three categories combined is also listed. It includes samples average, sample range and sample variability defined as twice the sample standard deviation. Variabilities smaller than 70 min are highlighted in blue ink. Starred societies mark studies conducted in the summer/spring season.

| Sleep timing (responses). Season: winter. |  |  |  |  |  |  |  |  |  |
| --- | --- | --- | --- | --- | --- | --- | --- | --- | --- |
|  |  | Offset |  |  | Midpoint |  | Onset |  |  |
|  |  | to noon | to WSR | WSA | to noon | to WSR | to noon | to WSR | to offset |
| | | $t_1$ | $t_1 - \beta D_w$ | $z_w$ | $t_2$ | $t_2 - \beta D_w$ | $t_3$ | $t_3 - \beta D_w - T$ | $t_3 - t_1 - T$ |
| *Industrial (N = 8)* |  |  |  |  |  |  |  |  |  |
| Denmark[11] | DNK | -5.2 h | -1.7 h | -13° | 2.9 h | 6.4 h | 11.0 h | -9.5 h | -7.8 h |
| Ireland[12] | IRL | -4.8 h | -1.1 h | -9° | 3.1 h | 6.8 h | 11.0 h | -9.3 h | -8.2 h |
| United Kingdom[13] | GBR | -4.8 h | -0.9 h | -8° | 3.0 h | 6.8 h | 10.7 h | -9.4 h | -8.5 h |
| France[14] | FRA | -5.8 h | -1.6 h | -15° | 2.1 h | 6.3 h | 10.0 h | -9.8 h | -8.3 h |
| Canada[17] | CAN | -5.3 h | -0.9 h | -9° | 2.6 h | 6.9 h | 10.4 h | -9.3 h | -8.3 h |
| Italy[15] | ITA | -5.0 h | -0.6 h | -6° | 2.8 h | 7.3 h | 10.6 h | -8.9 h | -8.3 h |
| Spain[16] | ESP | -5.5 h | -0.9 h | -10° | 2.5 h | 7.1 h | 10.5 h | -8.8 h | -7.9 h |
| United States[18] | USA | -5.6 h | -0.9 h | -10° | 2.3 h | 7.0 h | 10.2 h | -9.1 h | -8.2 h |
| *Pre-industrial (with electricity) (N = 5)* |  |  |  |  |  |  |  |  |  |
| Quilombolas[10] | QB2 | -4.9 h | +0.3 h | 2° | 2.8 h | 8.0 h | 10.6 h | -8.3 h | -8.6 h |
| Quilombolas[10] | QB1 | -5.5 h | -0.2 h | -3° | 2.4 h | 7.6 h | 10.2 h | -8.5 h | -8.3 h |
| Toba/Qom[8] | TQE | -5.2 h | +0.1 h | 1° | 2.2 h | 7.5 h | 9.6 h | -9.0 h | -9.2 h |
| Milange[9] | MIL | -5.7 h | -0.1 h | -3° | 1.9 h | 7.5 h | 9.5 h | -8.9 h | -8.8 h |
| *Chico Mendes[7] | CME | -7.1 h | -1.3 h | -19° | 1.2 h | 6.9 h | 9.4 h | -8.8 h | -7.5 h |
| *Pre-industrial (without electricity) (N = 9)* |  |  |  |  |  |  |  |  |  |
| Quilombolas[10] | QBN | -5.4 h | -0.1 h | -2° | 2.0 h | 7.3 h | 9.4 h | -9.3 h | -9.2 h |
| Toba/Qom[8] | TQN | -5.4 h | -0.0 h | -1° | 1.6 h | 7.0 h | 8.6 h | -10.0 h | -10.0 h |
| Fondwa[4] | FON | -6.9 h | -1.4 h | -20° | 1.1 h | 6.6 h | 9.1 h | -9.4 h | -8.0 h |
| Tengua[9] | TEN | -6.3 h | -0.7 h | -10° | 1.1 h | 6.7 h | 8.5 h | -9.9 h | -9.2 h |
| San[5] | SAN | -5.9 h | -0.3 h | -5° | 1.9 h | 7.5 h | 9.7 h | -8.7 h | -8.4 h |
| Tsimane[5] | TSI | -7.2 h | -1.6 h | -22° | 0.8 h | 6.4 h | 8.7 h | -9.7 h | -8.1 h |
| Mandena[5] | MAN | -6.1 h | -0.5 h | -8° | 0.8 h | 6.5 h | 7.8 h | -10.6 h | -10.1 h |
| *Chico Mendes[7] | CMN | -7.2 h | -1.5 h | -21° | 0.8 h | 6.6 h | 8.8 h | -9.5 h | -8.0 h |
| Hadza[5] | HAD | -6.9 h | -1.0 h | -14° | 1.2 h | 7.2 h | 9.4 h | -8.7 h | -7.7 h |
| *Univariate analysis for all three categories combined (N = 22)* |  |  |  |  |  |  |  |  |  |
| Sample average | $E(\{y_i\})$ | -5.8 h | -0.8 h | -9° | 2.0 h | 7.0 h | 9.7 h | -9.2 h | -8.5 h |
| Sample range | $\Delta\{y_i\}_{\min}^{\max}$ | 146 min | 119 min | 25° | 140 min | 101 min | 189 min | 136 min | 154 min |
| Sample variability | $2s(\{y_i\})$ | 96 min | 72 min | 14° | 92 min | 55 min | 106 min | 65 min | 82 min |

**Table III.** Sleep timing in industrial and in pre-industrial societies with and without electricity during the summer season. Times are expressed in decimal hours as a distance to solar noon *t*_*i*_ or a distance to winter sunrise (WSR) *t*_*i*_ − *βD_w_*. In the first case, the time offset (see Table I) and *T/*2 = 12 h must be added to obtain local times, one more hour must be added to industrial values due to DST. Summer solar altitude SSA at sleep offset and sleep onset distance to next sleep offset are also listed. Univariate analysis for all three categories combined is also listed. It includes samples average, sample range and sample variability defined as twice the sample standard deviation. Variabilities smaller than 70 min are highlighted in blue ink. Starred societies mark studies conducted in the autumn/winter season.

| Sleep timing (responses). Season: summer. |  |  |  |  |  |  |  |  |  |
| --- | --- | --- | --- | --- | --- | --- | --- | --- | --- |
|  |  | Offset |  |  | Midpoint |  | Onset |  |  |
|  |  | to noon | to WSR | SSA | to noon | to WSR | to noon | to WSR | to offset |
| | | $t_1$ | $t_1 - \beta D_w$ | $z_s$ | $t_2$ | $t_2 - \beta D_w$ | $t_3$ | $t_3 - \beta D_w - T$ | $t_3 - t_1 - T$ |
| *Industrial (N = 8)* |  |  |  |  |  |  |  |  |  |
| Denmark[11] | DNK | -6.2 h | -2.7 h | 18° | 1.9 h | 5.4 h | 10.0 h | -10.5 h | -7.8 h |
| Ireland[12] | IRL | -5.8 h | -2.1 h | 20° | 2.1 h | 5.8 h | 10.0 h | -10.3 h | -8.2 h |
| United Kingdom[13] | GBR | -5.8 h | -1.9 h | 20° | 2.0 h | 5.8 h | 9.7 h | -10.4 h | -8.5 h |
| France[14] | FRA | -6.8 h | -2.6 h | 10° | 1.1 h | 5.3 h | 9.0 h | -10.8 h | -8.3 h |
| Canada[17] | CAN | -6.3 h | -1.9 h | 14° | 1.6 h | 5.9 h | 9.4 h | -10.3 h | -8.3 h |
| Italy[15] | ITA | -6.0 h | -1.6 h | 16° | 1.8 h | 6.3 h | 9.6 h | -9.9 h | -8.3 h |
| Spain[16] | ESP | -6.5 h | -1.9 h | 9° | 1.5 h | 6.1 h | 9.5 h | -9.8 h | -7.9 h |
| United States[18] | USA | -6.6 h | -1.9 h | 7° | 1.3 h | 6.0 h | 9.2 h | -10.1 h | -8.2 h |
| *Pre-industrial (with electricity) (N = 3)* |  |  |  |  |  |  |  |  |  |
| Toba/Qom[8] | TQE | -6.2 h | -0.9 h | 6° | 1.8 h | 7.1 h | 9.9 h | -8.8 h | -7.9 h |
| *Milange[9] | MIL | -5.7 h | -0.1 h | 10° | 1.9 h | 7.5 h | 9.5 h | -8.9 h | -8.8 h |
| Chico Mendes[7] | CME | -6.9 h | -1.2 h | -8° | 1.3 h | 7.0 h | 9.5 h | -8.8 h | -7.6 h |
| *Pre-industrial (without electricity) (N = 8)* |  |  |  |  |  |  |  |  |  |
| Toba/Qom[8] | TQN | -6.3 h | -1.0 h | 5° | 1.2 h | 6.5 h | 8.7 h | -10.0 h | -9.0 h |
| *Fondwa[4] | FON | -6.9 h | -1.4 h | -5° | 1.1 h | 6.6 h | 9.1 h | -9.4 h | -8.0 h |
| *Tengua[9] | TEN | -6.3 h | -0.7 h | 3° | 1.1 h | 6.7 h | 8.5 h | -9.9 h | -9.2 h |
| San[5] | SAN | -5.6 h | +0.0 h | 11° | 2.7 h | 8.3 h | 10.9 h | -7.5 h | -7.5 h |
| Tsimane[5] | TSI | -7.9 h | -2.3 h | -19° | 0.6 h | 6.2 h | 9.0 h | -9.4 h | -7.1 h |
| *Mandena[5] | MAN | -6.1 h | -0.5 h | 4° | 0.8 h | 6.5 h | 7.8 h | -10.6 h | -10.1 h |
| Chico Mendes[7] | CMN | -7.1 h | -1.3 h | -10° | 0.9 h | 6.6 h | 8.8 h | -9.4 h | -8.1 h |
| Hadza[5] | HAD | -6.9 h | -1.0 h | -11° | 1.2 h | 7.2 h | 9.4 h | -8.7 h | -7.7 h |
| *Univariate analysis for all three categories combined (N = 19)* |  |  |  |  |  |  |  |  |  |
| Sample average | $E(\{y_i\})$ | -6.4 h | -1.4 h | 5° | 1.5 h | 6.5 h | 9.3 h | -9.7 h | -8.2 h |
| Sample range | $\Delta\{y_i\}_{\min}^{\max}$ | 137 min | 163 min | 39° | 126 min | 177 min | 186 min | 201 min | 178 min |
| Sample variability | $2s(\{y_i\})$ | 69 min | 93 min | 22° | 62 min | 88 min | 80 min | 101 min | 82 min |

**Table IV.** Multiple linear regression analyses for sleep timing in industrial and in pre-industrial societies. Predictor is always the shortest photoperiod *D*_*w*_(see Table I). Responses are listed in Table II (winter) and Table III (summer). Parentheses display a symmetric confidence semi-interval at the confidence level *α* = 5 %. Gradient and confidence interval are given in units of *β* = 30 min h^*−*1^, except starred values (WSA) which display units of degrees per hour. Gradients and confidence intervals for distance to WSR can be obtained adding 1 to numerical values listed for distance to noon. Probabilistic values smaller than the confidence level are highlighted in blue ink. Possible outliers are data whose confidence interval of residual at the confidence level excluded zero.

| Bivariate analysis: predictor is shortest photoperiod; responses are sleep timing. |  |  |  |  |  |  |
| --- | --- | --- | --- | --- | --- | --- |
| Response |  | Gradient | Confidence interval | Probabilistic value | Pearson's coefficient | Possible outliers |
| | $y$ | $m_s/\beta$ | $CI/\beta$ | $p$ | $R^2$ | |
| *Winter (N = 22)* |  |  |  |  |  |  |
| $t_1$, sleep offset | to noon | −0.77(37) | [−1.13, −0.40] | $< 10^{-3}$ | 0.49 | none |
|  | to winter sunrise |  |  | 0.2 | 0.082 | none |
| | winter solar altitude | −0.4(22)* | [−2.60, +1.86]* | 0.73 | $< 10^{-2}$ | none |
| $t_2$, midpoint | to noon | −0.85(28) | [−1.14, −0.57] | $< 10^{-5}$ | 0.66 | QB2 |
|  | to winter sunrise |  |  | 0.3 | 0.054 | QB2 |
| $t_3$, sleep onset | to noon | −0.95(35) | [−1.29, −0.60] | $< 10^{-4}$ | 0.62 | MAN |
| | to winter sunrise | | | 0.75 | $< 10^{-2}$ | MAN |
|  | to sleep offset | −0.18(43) | [−0.61, +0.24] | 0.38 | 0.038 | TQN MAN |
| *Summer (N = 19)* |  |  |  |  |  |  |
| $t_1$, sleep offset | to noon | −0.27(34) | [−0.62, +0.07] | 0.11 | 0.14 | TSI |
| | to winter sunrise | | | $< 10^{-3}$ | 0.54 | TSI |
| $t_2$, midpoint | to noon | −0.28(30) | [−0.58, +0.02] | 0.069 | 0.18 | SAN |
| | to winter sunrise | | | $< 10^{-3}$ | 0.59 | SAN |
| $t_3$, sleep onset | to noon | −0.28(41) | [−0.69, +0.12] | 0.16 | 0.11 | SAN MAN |
| | to winter sunrise | | | $< 10^{-2}$ | 0.45 | SAN MAN |
| | to sleep offset | −0.09(44) | [−0.45, +0.43] | 0.97 | $< 10^{-3}$ | MAN |

## References

[1] Borbély, A. A. A two process model of sleep regulation. Hum. Neurobiol. 1, 195–204 (1982). DOI 10.1111/jsr.12371.

[2] Ekirch, A. R. Sleep we have lost: Pre-industrial slumber in the British Isles. Am. Hist. Rev. 106, 343–386 (2001).

[3] Bin, Y. S., Marshall, N. S. & Glozier, N. Secular trends in adult sleep duration: A systematic review. Sleep Med. Rev. 16, 223–230 (2012). URL http://linkinghub.elsevier.com/retrieve/pii/S1087079211000736. DOI 10.1016/j.smrv.2011.07.003.

[4] Knutson, K. L. Sleep duration, quality, and timing and their associations with age in a community with-out electricity in Haiti. Am. J. Hum. Biol. 26, 80–86 (2014). URL http://doi.wiley.com/10.1002/ajhb.22481. DOI 10.1002/ajhb.22481.

[5] Yetish, G. et al. Natural sleep and its seasonal variations in three pre-industrial societies. Curr. Biol. 25, 2862–2868 (2015). URL https://www.cell.com/current-biology/comments/S0960-9822(15)01157-4. DOI 10.1016/j.cub.2015.09.046.

[6] Samson, D. R. et al. Segmented sleep in a non-electric, small-scale agricultural society in Madagascar. Am. J. Hum. Biol. 29, e22979 (2017). URL http://www.ncbi.nlm.nih.gov/pubmed/28181718. DOI 10.1002/ajhb.22979.

[7] Moreno, C. R. C. et al. Sleep patterns in Amazon rubber tappers with and without electric light at home. Sci. Rep. 5, 14074 (2015). URL http://www.nature.com/articles/srep14074. DOI 10.1038/srep14074.

[8] de la Iglesia, H. O. et al. Access to electric light is associated with shorter sleep duration in a traditionally hunter-gatherer community. J. Biol. Rhythms 30, 342–350 (2015). DOI 10.1177/0748730415590702.

[9] Beale, A. D. et al. Comparison between an African town and a neighbouring village shows delayed, but not decreased, sleep during the early stages of urbanisation. Sci. Rep. 7, 5697 (2017). URL http://www.nature.com/articles/s41598-017-05712-3. DOI 10.1038/s41598-017-05712-3.

[10] Pilz, L. K., Levandovski, R., Oliveira, M. A. B., Hi-dalgo, M. P. & Roenneberg, T. Sleep and light exposure across different levels of urbanisation in Brazilian communities. Sci. Rep. 8, 11389 (2018). URL http://www.nature.com/articles/s41598-018-29494-4. DOI 10.1038/s41598-018-29494-4.

[11] Det Nationale Forkskningscenter for Velfœrd. Danish Time Use Survey: Danske Tidsanvendelseundersøgelsen. Center for Survey and Survey/Register Data (distribuitor) (2001). URL http://bit.ly/2t37Ksj.

[12] Economic and Social Research Institute. The Irish National Time-Use Survey. Irish Social Sience Data Archive (distribuitor) (2005).

[13] Ipsos-RSL and Office of National Statistics. United Kingdom Time Use Survey 2000 (computer file). 3rd ed, Colchester, Essex: UK Data archive (distribuitor) (2003).

[14] L’Institut National de la Statisque et des études économiques. French Time Use Survey. Enquäte emploi du Temps et Décisions dans les couples (2010).

[15] L’Istituto nazionale di statistica (Istat). Italian Time Use Survey: Uso del tempo (2009).

[16] Instituto Nacional de Estadística. Spanish Time Use Survey: Encuesta de Empleo del Tiempo (2010). URL http://bit.ly/2uQFqa0.

[17] Statistics Canada/Statisque Canada. General Social Survey, Time Use. cycle 19. computer file (2005).

[18] Bureau of Labor Statistics. American Time Use Survey. computer file (multi year data) (2012). URL https://www.bls.gov/tus/datafiles_2013.htm.

[19] Martín-Olalla, J. M. Latitudinal trends in human primary activities: characterizing the winter day as a synchronizer. Sci. Reports 2018 81 8, 5350 (2018). URL https://www.nature.com/articles/s41598-018-23546-5. DOI 10.1038/s41598-018-23546-5.

[20] Fisher, S. P., Foster, R. G. & Peirson, S. N. The Circadian Control of Sleep. In Circadian Clocks, 157–183 (Springer, Berlin, Heidelberg, 2013). URL http://link.springer.com/10.1007/978-3-642-25950-0_7. DOI 10.1007/978-3-642-25950-0_7.

[21] Borbély, A. A., Daan, S., Wirz-Justice, A. & Deboer, T. The two-process model of sleep regulation: a reappraisal. J. Sleep Res. 25, 131–143 (2016). URL http://doi.wiley.com/10.1111/jsr.12371. DOI 10.1111/jsr.12371.

[22] Ekirch, A. R. At day’s close : night in times past (Norton, 2005).

[23] Ekirch, A. R. Segmented Sleep in Preindustrial Societies. Sleep 39, 715–716 (2016). URL http://www.journalsleep.org/ViewAbstract.aspx?pid=30498. DOI 10.5665/sleep.5558.

[24] Kantermann, T., Juda, M., Merrow, M. & Roenneberg, T. The Human Circadian Clock’s Seasonal Adjustment Is Disrupted by Daylight Saving Time. Curr. Biol. 17, 1996–2000 (2007). URL https://www.cell.com/abstract/S0960-9822(07)02086-6. DOI 10.1016/j.cub.2007.10.025.

[25] Bartky, I. R. One Time Fits All (Standford University Press, 2007).

[26] Roenneberg, T. & Merrow, M. The Circadian Clock and Human Health. Curr. Biol. 26, R432–43 (2016). URL http://www.ncbi.nlm.nih.gov/pubmed/27218855. DOI 10.1016/j.cub.2016.04.011.

[27] Meira e Cruz, M. et al. Impact of Daylight Saving Time on circadian timing system: An expert statement. Eur. J. Intern. Med. 3–5 (2019). URL https://www.sciencedirect.com/science/article/pii/S0953620519300135. DOI 10.1016/j.ejim.2019.01.001.

